# Neisseria gonorrhoeae induces local secretion of IL-10 at the human cervix to promote asymptomatic colonization

**DOI:** 10.1101/2024.05.23.595602

**Authors:** Yiwei Dai, Qian Yu, Vonetta Edwards, Hervé Tettelin, Daniel C. Stein, Wenxia Song

## Abstract

Gonorrhea, caused by the human-restricted pathogen *Neisseria gonorrhoeae*, is a commonly reported sexually transmitted infection. Since most infections in women are asymptomatic, the true number of infections is likely much higher than reported. How gonococci (GC) colonize women’s vaginocervix without triggering symptoms remains elusive. Using a human cervical tissue explant model, we found that GC inoculation increased the local secretion of both pro- (IL-1β and TNF-α) and anti-inflammatory (IL-10) cytokines during the first 24-h. Cytokine induction required GC expression of an Opa isoform that binds the host receptors carcinoembryonic antigen-related cell adhesion molecules (CEACAMs). GC inoculation induced NF-κB activation in both cervical epithelial and subepithelial cells. However, inhibition of NF-κB activation, which reduced GC-induced IL-1β and TNF-α, did not affect GC colonization. Neutralizing IL-10 or blocking IL-10 receptors by antibodies reduced GC colonization by increasing epithelial shedding and epithelial cell-cell junction disassembly. Inhibition of the CEACAM downstream signaling molecule SHP1/2, which reduced GC colonization and increased epithelial shedding, reduced GC-induced IL-10 secretion. These results show that GC induce local IL-10 secretion at the cervix by engaging the host CEACAMs to prevent GC-colonizing epithelial cells from shedding while suppressing inflammation activation, providing a potential mechanism for GC asymptomatic infection in women.

## Introduction

Gonorrhea, caused by a Gram-negative bacterium, *Neisseria gonorrhoeae,* is the second most common sexually transmitted infection, with 648,056 cases reported in the US in 2022 (1). In women, most GC infections are asymptomatic or subclinical; therefore, the true number of infections is likely far higher than reported (2, 3). Gonococci (GC) infect humans exclusively, primarily targeting male and female urethras and the female vaginocervix. GC can ascend from the vagina to the upper female reproductive tract (FRT) through the cervix (4), leading to severe clinical consequences, including pelvic inflammatory disease and infertility (5, 6). Asymptomatic infections allow bacteria to silently spread among sex partners and delay the treatment until the infection permanently damages the FRT. How GC colonize the human vaginocervix without inducing symptoms or eliciting immune responses remains elusive.

GC initiate infection at the FRT by colonizing the luminal epithelial cells of the vaginocervix (4, 7). GC can invade into epithelial cells (8, 9) and transmigrate across columnar epithelial cells (10, 11), the latter of which is associated with disseminated gonococcal infections (12). GC initiate the interaction with epithelial cells using pili (13, 14). Pili retraction brings GC close to the luminal surface of epithelial cells (14, 15), which allows the interaction of GC surface molecules, such as opacity-associated proteins (Opa), with their human-specific host receptors, such as carcinoembryonic antigen-related cell adhesion molecules (CEACAMs) and heparan sulfate proteoglycans (HSPGs) (16, 17). Using a human cervical tissue explant model, we have shown that GC preferentially colonize the ectocervix and transformation zone but selectively penetrate into the subepithelia of the transformation zone and endocervix, mimicking observations of patients’ biopsies (4, 7, 18). Pili are essential for GC colonization at the cervical epithelia, including the multi-layer squamous ectocervical and single-layered columnar endocervical epithelial cells (18). The expression of HSPG-binding or no Opas induces cervical epithelial cell shedding by disrupting epithelial cell-cell junction, a mechanism used by the host to eliminate colonizing GC (11, 18–20). In contrast, the expression of Opas binding to CEACAMs (Opa_CEA_) promotes GC colonization at the human cervix by inhibiting GC-induced epithelial cell-cell junction disruption and epithelial shedding (18). The inhibitory effects of Opa_CEA_ on cervical colonization depend on the immunoreceptor tyrosine-based inhibitory motif (ITIM) in the cytoplasmic tails of CEACAM1 and its downstream signaling molecule SH2-containing protein tyrosine phosphatase (SHP) 1/2 (18). GC-induced epithelial cell shedding and the inhibitory effects of Opa_CEA_ have also been observed in the vaginocervix of vaginally infected mice that express human CEACAM5 and in human epithelial cell lines (19, 20). GC-induced epithelial cell-cell junction disruption enables GC to penetrate through the endocervical and transformation zone epithelia and enter subepithelial tissues. Opa_CEA_ inhibits GC penetration by restoring epithelial cell-cell junction complexes (11, 18). Thus, GC can switch their infection strategies by phase variation of their Opa proteins targeting different host receptors. The fact that most of the 11 isoforms of Opa proteins bind to CEACAMs (21, 22) suggests that Opa_CEA_-promoted colonization but not penetration enables GC survival on the human cervix.

Bacterial colonization of the epithelium can induce inflammatory chemokine and cytokine production as warning signals by activating receptors recognizing pathogen-associated molecular patterns on the epithelial cell surface (23–25). Inflammatory chemokines and cytokines can chemoattract and activate resident and circulating immune cells to clear the infection. Neutrophil infiltration is the main clinical feature of symptomatic GC infection (26). How GC escape or suppress such immune detection to colonize women’s vaginocervix silently remains a critically unanswered question. The relationship between GC asymptomatic infection and the initial local cytokine response of the vaginocervix is unknown. Both inflammatory cytokines, such as IL-1β, TNF-α, and IL-17, and anti-inflammatory cytokines, such as IL-10 and TGF-β, have been detected in the cervical mucus of uninfected women (27–29). The elevation, reduction, and no change of both pro- and anti-inflammatory cytokines have all been reported previously in the vaginocervical secretion of patients with GC infection, compared to uninfected controls (27, 30–33). This broad spectrum of data probably resulted from variations in sample collecting methods, sample collecting times relative to the menstrual cycle and GC infection course, and whether the infection was symptomatic and asymptomatic. In vitro studies have shown that GC induce the production of the anti-inflammatory cytokines IL-10 by human monocyte-derived dendritic cells (34), mouse bone marrow-derived dendritic cells (35), human monocyte-derived macrophages (36), human CD4+ T cells (37), human and mouse peripheral blood mononuclear cells (38, 39), mouse genital tract tissue explants (39), and cells from mouse iliac lymph nodes (39). As IL-10 is a master regulator of immunity and functions to protect hosts from over-exuberant responses (40–42), these data suggest inducing an anti-inflammatory cytokine response is a potential mechanism for GC to evade the host adaptive immunity when GC can enter tissues and directly interact with immune cells. However, the kind of cytokine responses colonizing GC can induce initially and locally at the human cervix and how induced cytokines impact immune detection are unknown.

While the epithelium can initiate the local cytokine responses upon pathogen interaction and pathogen-induced wounding, cytokines have been shown to regulate the barrier function and the homeostasis of the epithelium (43–45), potentially influencing the infection processes. The pro-inflammatory cytokines IL-1β and TNF-α, which are associated with inflammatory bowel diseases, disrupt epithelial cell-cell junctions by increasing the expression and activation levels of myosin light chain kinase (MLCK) (46–48). TNF-α can also damage the epithelial barrier by inducing epithelial apoptosis, leading to shedding (49). In contrast, the anti-inflammatory cytokine IL-10 activates epithelial cell proliferation and wound repair, strengthening their barrier function (41, 42, 50, 51). IL-10 or IL-10 receptor (IL-10R) deficiency leads to human inflammatory bowel diseases (52, 53). However, the pleiotropic cytokine TGF-β can strengthen (such as in the intestine) (54) or weaken (such as in the uterus) (55) epithelial cell-cell junctions depending on its anatomic location. The cytokine environment of the human cervix and its role in GC infection have not been well studied.

This study examined the relationship between locally secreted cytokines and GC infection when the cervix is initially exposed to GC, using our previously established human cervical tissue explant model, which mimics *in vivo* GC infection and eliminates the influence of blood and lymphoid circulation. We focused on cytokines detected in female secretions and known to regulate epithelial integrity. Our results show that GC inoculation increases the secretion of the pro-inflammatory cytokines TNF-α and IL-1β by activating NF-κB and the secretion of the anti-inflammatory cytokine IL-10 by engaging the host receptor CEACAMs by Opa and activating CEACAM1 downstream signaling molecule SHP1/2. GC-induced pro-inflammatory cytokines TNF-α and IL-1β do not significantly affect GC colonization at the ectocervix. However, GC-induced anti-inflammatory cytokine IL-10 enhances GC colonization at the ectocervix by inhibiting GC-induced epithelial cell-cell junction disruption and the shedding of GC-associated cervical epithelial cells.

## Results

### GC inoculation increases the local secretion levels of both pro- and anti-inflammatory cytokines in the human cervix

To determine the initial, local cytokine response of the cervix to GC infection, we utilized the human cervical tissue explant model, as it mimics GC infection as observed *in vivo* using patients’ biopsies (4, 7, 11, 18) and is disconnected from the blood and lymphatic circulation. Human cervical tissue explants (ecto- to endocervix) from the same donor were incubated without (No GC) or with MS11 GC strains with either all 11 isoforms of *opa* genes deleted (MS11ΔOpa) or the expression of a single non-phase variable Opa_52_ protein that binds to CEACAMs (MS11Opa_CEA_) at a MOI of 10 (10 bacteria per luminal epithelial cell) for 24 h at 37°C. Culture supernatants were collected to quantify cytokine secretion using ELISA and Luminex Magpix. Cytokine levels were normalized to the luminal surface area of tissue explants and the volume of supernatant. Uninoculated tissue explants secreted detectable or modest amounts of pro-inflammatory cytokines IL-1β (Figure 1A) and TNF-α (Figure 1B), the anti-inflammatory cytokine IL-10 (Figure 1C), and the pleiotropic cytokine TGF-β (Figure 1D). Inoculation with MS11Opa_CEA_, which colonize the cervical epithelium more efficiently than MS11ΔOpa (18), significantly increased both the pro- and anti-inflammatory cytokines but not TGF-β (Figure 1, A-D). Inoculation with MS11ΔOpa, which colonize less but penetrate more efficiently than MS11Opa_CEA_ (18), only moderately elevated the level of IL-1β (Figure 1A) but not TNF-α, IL-10, and TGF-β (Figure 1, B-D). IL-17 was undetectable in culture supernatants of GC uninoculated and inoculated cervical tissue explants. When inoculating explants containing the ectocervix alone, MS11Opa_CEA_ induced similar increases in IL-1β (Figure 1E), TNF-α (Figure 1F), and IL-10 (Figure 1G) as inoculated explants containing all three cervical regions. These data suggest that GC infection elevates the local secretion levels of both pro- and anti-inflammatory cytokines by the cervix during the first 24 h in an Opa_CEA_-dependent manner.

**Figure 1.**
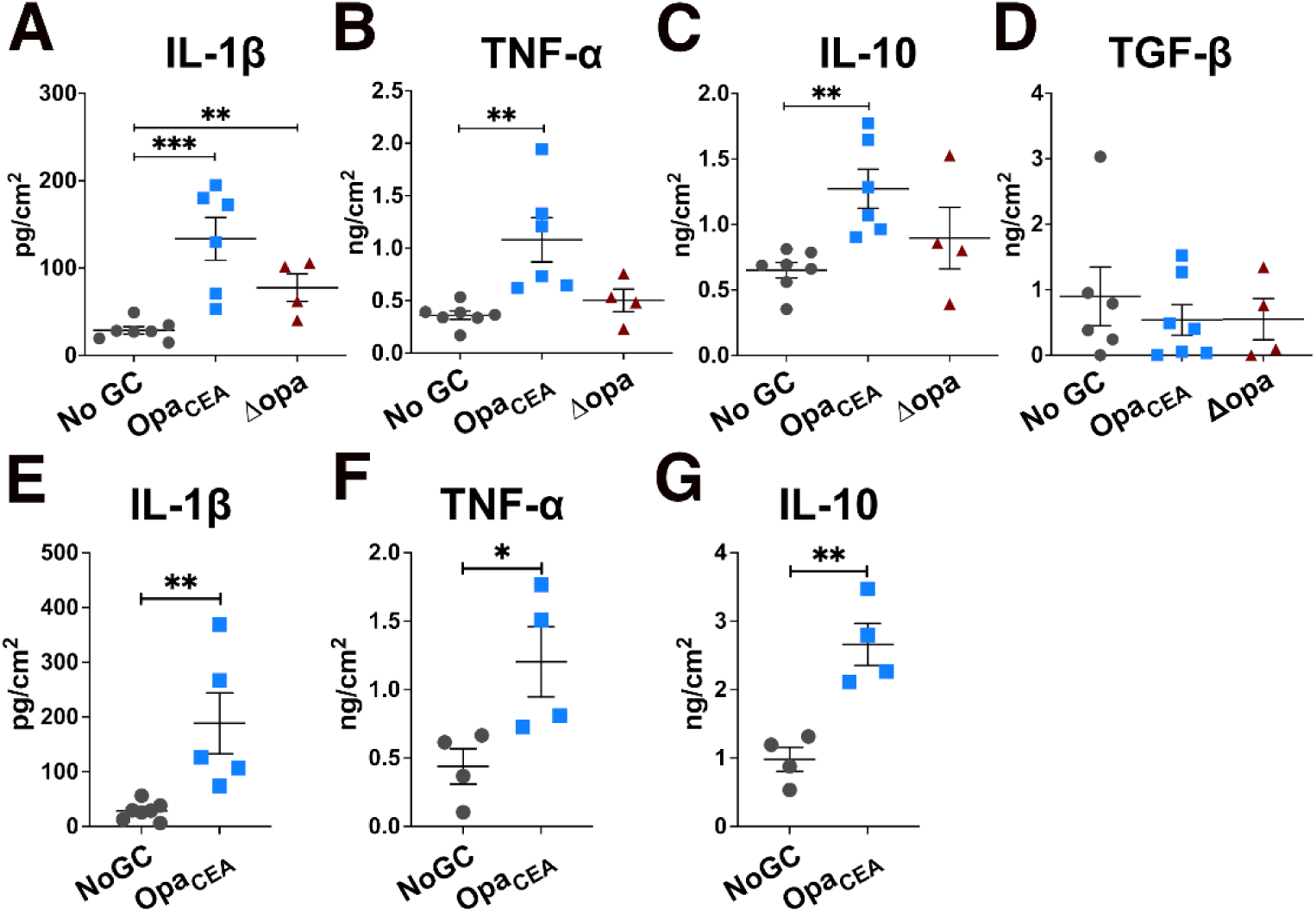
GC inoculation increases the secretion of both pro- and anti-inflammatory cytokines by human cervical tissue explants. Human cervical tissue explants with the ecto- to endocervix (**A-D**) or with the ectocervix alone (**E-G**) were incubated without (No GC) or with MS11 GC strains expressing no *opa* genes (ΔOpa) or a non-phase variable Opa_52_ that binds to CEACAMs (Opa_CEA_) at a MOI of 10 for 24 h at 37°C. Supernatants of cervical tissue explant media were collected. The average concentrations (±SEM) of IL-1β (**A, E**), TNF-α (**B, F**), IL-10 (**C, G**), and TGF-β (**D**) were measured by Luminex Magpix (IL-1β, TNF-α, and IL-10) or ELISA (TGF-β), normalized to the luminal surface areas of tissue explants and supernatant volumes. Data points represent cervical tissues from individual human subjects. The data were generated from 4-7 cervixes. * *p*<0.05, ** *p*<0.01, *** *p*<0.001, by unpaired student’s *t*-test.

### GC-induced pro-inflammatory cytokine secretion in the human cervix depends on NF-κB

The transcription factor NF-κB is a crucial regulator of cytokine production (56, 57). To examine the possible involvement of NF-κB in GC-mediated cytokine expression, we determined if GC infection activated NF-κB in cervical cells using immunofluorescence microscopy. We measured the mean fluorescence intensity (MFI) of the NF-κB p65 subunit in the nuclei of individual cervical cells within the epithelium and subepithelium (135 μm below the epithelium) (Figure 2A). MS11Opa_CEA_ increased NF-κB p65 MFI in the nuclei of both epithelial and subepithelial cells of all three cervical regions, ectocervix (Ecto), transformation zone (TZ), and endocervix (Endo), compared to no GC controls (Figure 2B). Among the three cervical regions of MS11Opa_CEA_-infected tissue explants, the increases in the nuclear NF-κB p65 MFI of the ectocervical epithelial and subepithelial cells were higher than the other two cervical regions (Figure 2B). However, MS11ΔOpa did not significantly change the NF-κB p65 MFI of the cervical epithelial cell nuclei (Figure 2B, left panel). Interestingly, MS11ΔOpa reduced the nuclear NF-κB p65 MFI of subepithelial cells in the two GC-penetrated cervical regions, while slightly increasing the nuclear NF-κB p65 MFI of ectocervical subepithelial cells where MS11ΔOpa cannot penetrate (Figure 2B, right panel). Furthermore, both strains significantly increased the mRNA expression levels of the NF-κB genes NFKB2 and/or NFKB1 and genes encoding proteins that act upstream of NF-κB activation and downstream of TNF-α and IL-1β, TRAF1, TRAF3, TRAF4, IRAK2, and/or IRAK3, as measured by whole tissue NanoString-based transcriptomic analysis (Figure 2C). These data indicate that MS11Opa_CEA_ inoculation increases NF-κB activation levels of both cervical epithelial and subepithelial cells.

**Figure 2.**
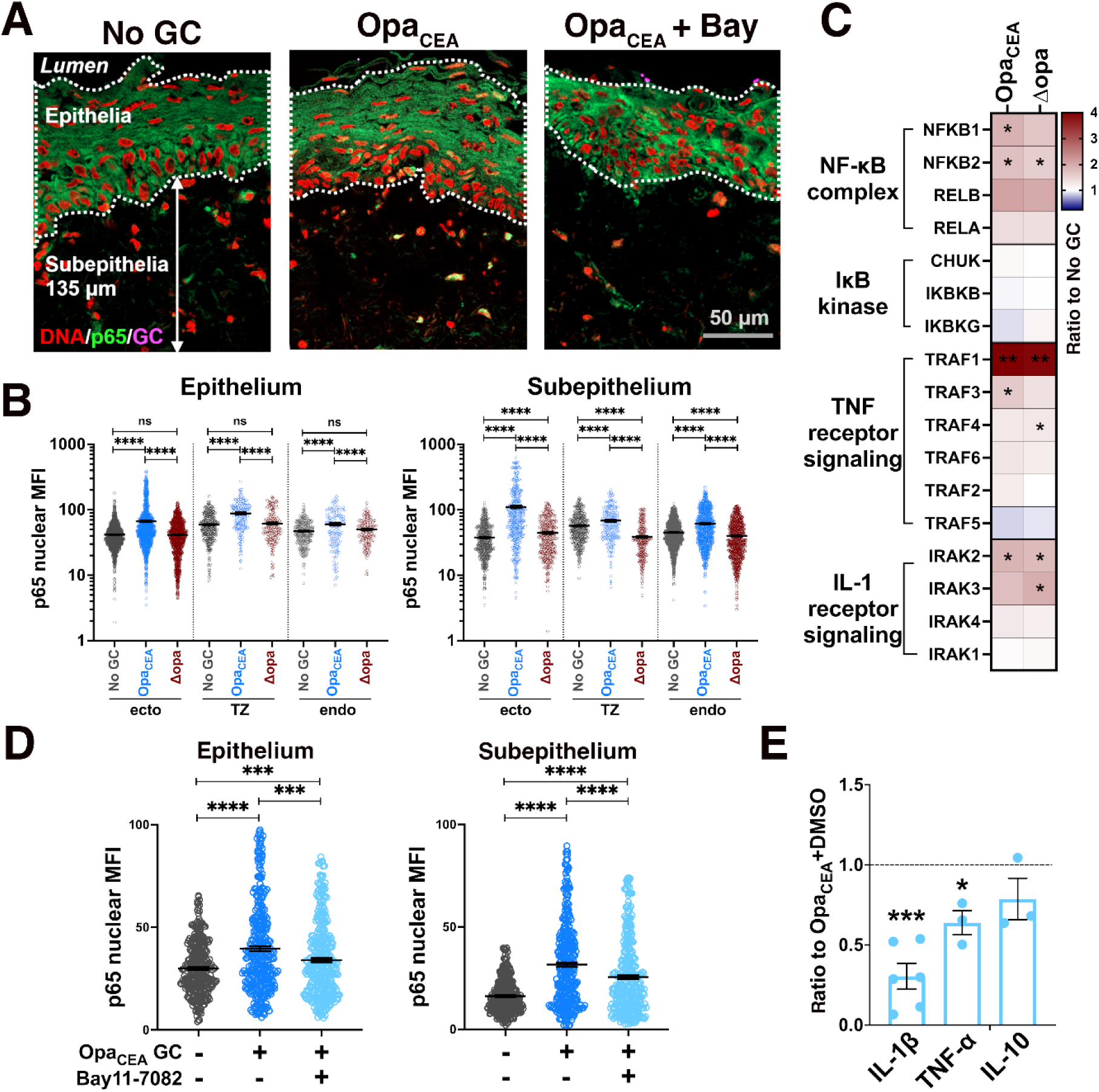
NF-κB is involved in GC-induced pro-inflammatory cytokine production by the human cervix. Human cervical tissue explants were incubated without or with MS11Opa_CEA_ or MS11ΔOpa (MOI~10) in the absence or presence of the NF-κB inhibitor Bay11-7082 (3 μM) for 24 h. The culture supernatants were collected, and tissues were cryopreserved. (**A**) Tissue sections were stained for NF-κB p65 and GC by antibodies and nuclei by Hoechst and imaged using a confocal fluorescence microscope (CFM). Representative images of the ectocervix without or with MS11Opa_CEA_ GC inoculation and the NF-κB inhibitor Bay11-7082 are shown. White dished lines outline the epithelium, and arrows indicate the 135 μm depth of the subepithelium. Scale bar, 50 μm. (**B**) The mean fluorescent intensity (MFI) (±SEM) of NF-κB p65 in the nuclei of individual epithelial and subepithelial cells (135 μm below the epithelium) in the ectocervix, transformation zone, and endocervix were measured. Data points represent individual nuclei. Data were generated from cervical tissues from 3-4 human subjects, 2 independent analyses per cervix, and 35-200 epithelial and 45-170 subepithelial cells per analysis per cervical region. (**C**) A heatmap shows the relative mRNA levels of NF-κB-related genes determined by the NanoString Immunology panel. RNAs were isolated from cervical tissue explants inoculated without (no GC) or with MS11Opa_CEA_ or ΔOpa GC. The comparisons of MS11Opa_CEA_- or ΔOpa-inoculated with no GC control tissues from 3 human subjects are shown, and stars indicate *p* values by unpaired student’s *t*-test. (**D**) The nuclear MFI of NF-κB p65 in individual epithelial and subepithelial cells from the ectocervical tissue explants inoculated without or with MS11Opa_CEA_ in the absence or presence of Bay11-7082. Datapoint represents individual nuclei. The average values (±SEM) were generated from ectocervical tissues from 3 human subjects, 3 independent analyses per cervix, and 40 nuclei per region per analysis. (**E**) The concentrations of IL-1β, TNF-α, and IL-10 in the supernatants of ectocervical tissue explants inoculated with MS11Opa_CEA_ in the absence and presence of Bay11-7082 were measured using Luminex Magpix (IL-1β, TNF-α and IL-10) and ELISA (IL-1β) and normalized to the luminal surface areas and the supernatant volume of individual explants. Shown are the ratios of cytokine concentrations (±SEM) secreted by ectocervical tissue explants inoculated with MS11Opa_CEA_ in the presence of Bay11-7082 compared to those MS11Opa_CEA_ inoculated in the presence of the vehicle control DMSO. Data points represent individual analysis. * *p*<0.05, ** *p*<0.01, *** *p*<0.001, **** *p*<0.0001, by unpaired student’s *t*-test.

To determine if GC-induced NF-κB activation is involved in the cytokine induction, we utilized a membrane-permeable small molecule inhibitor of NF-κB, Bay11-7082, that inhibits IκBα phosphorylation (58). The inhibitor (3 µM) was included in the 24 h incubation of ectocervical tissue explants with MS11Opa_CEA_. Inhibitor treatment significantly reduced the NF-κB p65 MFI in the nuclei of ectocervical epithelial and subepithelial cells, even though the treatment did not bring their nuclear NF-κB p65 MFI back to the no GC level (Figure 2D). Notably, the inhibitor treatment significantly reduced the secretion levels of the inflammatory cytokines IL-1β and TNF-α in the ectocervix but did not have a significant impact on the ectocervical secretion of the anti-inflammatory cytokine IL-10 (Figure 2E). These data suggest that GC-induced pro- but not anti-inflammatory cytokine secretion involves NF-κB.

### Reductions in pro-inflammatory cytokines by NF-κB inhibition do not impact GC colonization of the ectocervix

Treatment with the pro-inflammatory cytokine IL-1β or TNF-α disrupts intestinal epithelial cell-cell junctions and induces epithelial cell apoptosis, in addition to inducing inflammation (43, 46). We determined whether NF-κB inhibition, which reduced IL-1β and TNF-α production (Figure 2E), affected GC infection. Ectocervical tissue explants inoculated with MS11Opa_CEA_ in the absence or presence of the NF-κB inhibitory Bay11-7082 (3 μM) were stained for GC (antibodies), DNA (Hoechst) to mark individual cells, and F-actin (phalloidin) to mark each ectocervical cell (Figure 3A). We quantified GC colonization by measuring the percentage of luminal epithelial cells with attached GC (Figure 3A, left panel), reflecting the extent of the epithelium being infected and GC fluorescence intensity (FI) per μm^2^ of the luminal surface (Figure 3A, right panel), reflecting the relative amount of GC colonizing the ectocervical epithelium. Treatment with the NF-κB inhibitor changed neither the percentage of luminal ectocervical epithelial cells associated with GC (Figure 3B, left panel) nor the GC FI per μm^2^ of the luminal surface significantly (Figure 3B, right panel). The NF-κB inhibitor also did not affect GC growth (Supplementary Figure 1). We further quantified ectocervical epithelial cell shedding by measuring the percentage of the epithelial thickness and the percentage of epithelial cell layers remaining using no GC controls as 100% (Figure 3C). NF-κB inhibitor treatment did not affect the thickness (Figure 3D, left panel) or the cell layer number (Figure 3D, right panel) of the ectocervical epithelium. Even though the secreting levels of IL1β and TNF-α were elevated in the absence of the NF-κB inhibitor (Fig 1A and B), MS11Opa_CEA_ GC inoculation did not induce significant changes in epithelial thickness and cell layers (Figure 3D). Together, these data suggest that GC-induced pro-inflammatory cytokines do not interfere with GC colonization of the human cervix.

**Figure 3.**
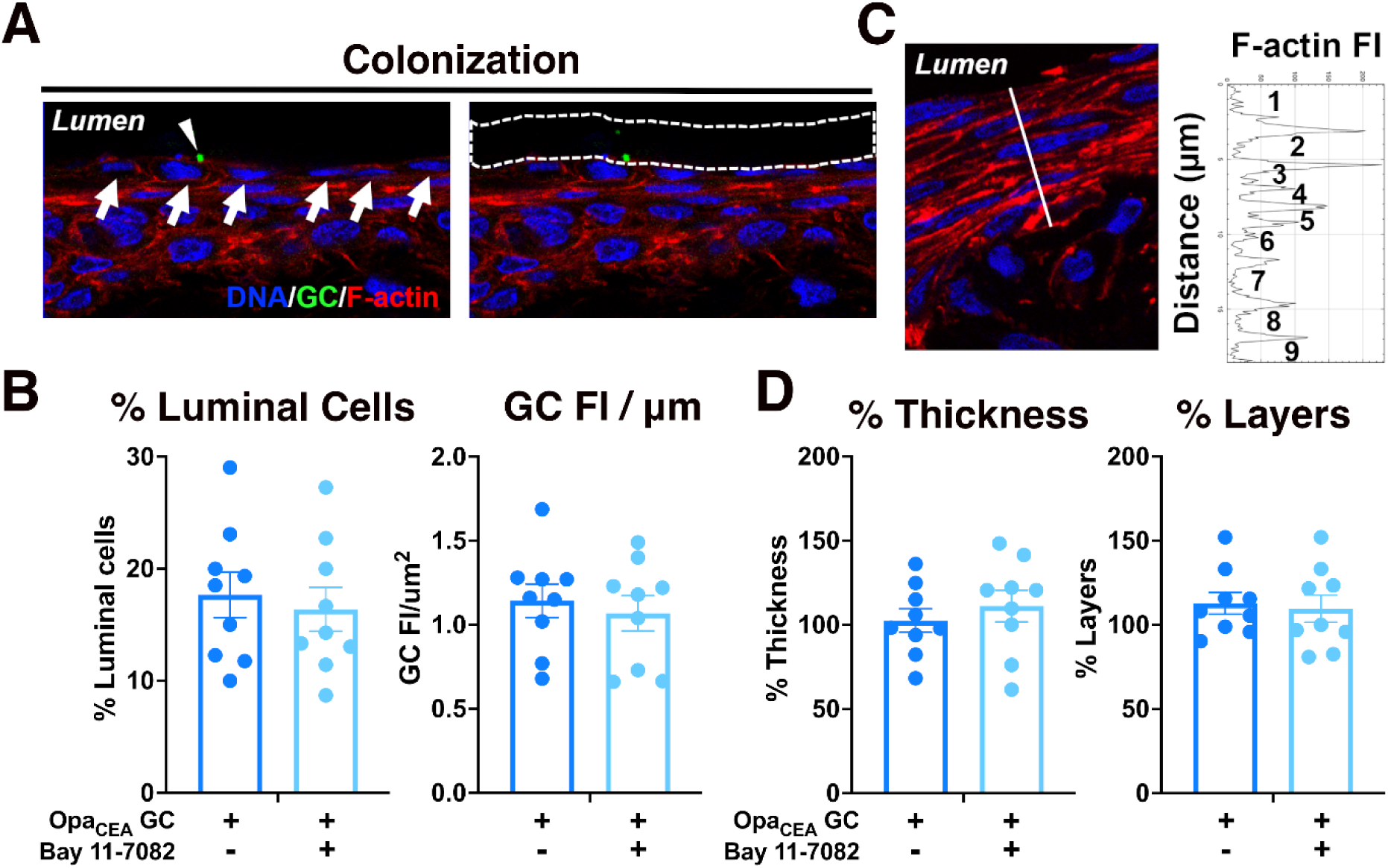
Pro-inflammatory cytokine reduction by NF-κB inhibition does not affect GC colonization of the ectocervix and ectocervical epithelial shedding. Human ectocervical tissue explants were inoculated without or with MS11Opa_CEA_ in the absence and presence of the NF-κB inhibitor Bay11-7082 (3 μM) for 24 h (MOI~10) and cryopreserved. Tissue sections were stained for GC by a polyclonal antibody, DNA by Hoechst, and F-actin by phalloidin and analyzed using CFM. (**A**) Quantification of GC colonization by the percentage (±SEM) of luminal epithelial cells associated with GC (left panel) and by fluorescence intensity (FI) (±SEM) of GC staining per μm^2^ of the luminal surface (right panel). Arrows in the left panel point to individual luminal epithelial cells and arrowhead to a GC-associated luminal epithelial cell. White dashed lines in the right panel outline the luminal surface area where GC FI was measured. (**B**) The average percentages (±SEM) of luminal epithelial cells associated with GC and the average value of GC FI per μm^2^ of the luminal surface from ectocervical tissues of 3 human subjects with 3 independent analyses per cervix and 3-10 randomly acquired images per analysis. Data points represent each independent analysis. (**C**) Quantification of epithelium shedding by the percentage of remaining epithelium thickness (left panel, the length of the dashed line vertical against the epithelium) and cell layers based on the FI line profile of F-actin staining on the dashed line (right panel), compared with no GC control (**C**, right). (**D**) The average percentages (±SEM) of the remaining epithelial thickness and cell layers from ectocervical tissues of 3 human subjects with 3 independent analyses per cervix, 3-10 randomly acquired images per analysis, and 3 line profiles per image. Data points represent each independent analysis. There are no significant differences by unpaired student’s *t*-test.

### The anti-inflammatory cytokine IL-10 promotes GC colonization by inhibiting epithelial cell shedding

In contrast to the pro-inflammatory cytokines IL-1β and TNF-α, the anti-inflammatory cytokine IL-10 has been shown to strengthen the barrier and wound repair functions of the intestinal epithelium (43, 50, 51). We determined whether MS11Opa_CEA_-induced IL-10 production contributes to GC infection, using either an IL-10 or IL-10 receptor (IL-10R) α antibody to prevent secreted IL-10 from engaging IL-10R (59, 60). Ectocervical tissue explants were inoculated with MS11Opa_CEA_ in the absence or presence of IL-10 (10 µg/ml) or IL-10Rα (5 µg/ml) antibodies for 24 h. Luminex Magpix and ELISA analysis found that the IL-10 antibody effectively reduced the IL-10 level but did not significantly affect the levels of the pro-inflammatory cytokines IL-1β and TNF-α in the supernatants of ectocervical tissue explants inoculated with MS11Opa_CEA_ (Figure 4A). IL-10 and IL-10Rα antibodies did not directly affect GC growth (Supplementary Figure 2). Immunofluorescence microscopic analysis found that the IL-10Rα antibody primarily concentrated at the ectocervical epithelium but could also be detected in the subepithelium (Supplementary Figure 3). Importantly, both IL-10 and IL-10Rα antibodies significantly reduced the percentages of the luminal epithelial cells associated with GC (Figure 4, B and C) and the GC FI per µm^2^ (Figure 4, B and D), indicating reductions in GC colonization. Increased disassociation of GC-attached epithelial cells from the ectocervix was clearly visible in immunofluorescence images of IL-10 or IL-10Rα antibody-treated tissue explants, compared to untreated tissue explants (Figure 4B, arrowheads). We quantified epithelial cell shedding using CFM images. Treatment with IL-10 and IL-10Rα antibodies reduced epithelial remaining thickness (Figure 4E) and cell layer number (Figure 4F), as compared to untreated MS11Opa_CEA_-infected ectocervical tissue explants as controls, indicating increased epithelial cell shedding. Using CFM images, we explored if IL-10 neutralization or IL-10Rα blocking induced epithelial shedding by disrupting epithelial cell-cell junction. We measured the fluorescence intensity ratios (FIR) of the ectocervical epithelial cell-cell junctional protein E-cadherin (E-cad) at the cell-cell junction relative to the cytoplasm using FI line profiles that vertically cross the epithelial cell-cell contact surface (Figure 4G). Treatment of IL-10 or IL-10Rα antibodies significantly reduced the junction to cytoplasm ratios of E-cad FI (Figure 4H). Our results suggest that MS11Opa_CEA_-induced IL-10 promotes bacterial colonization by strengthening epithelial cell-cell junctions and preventing epithelial cell shedding.

**Figure 4.**
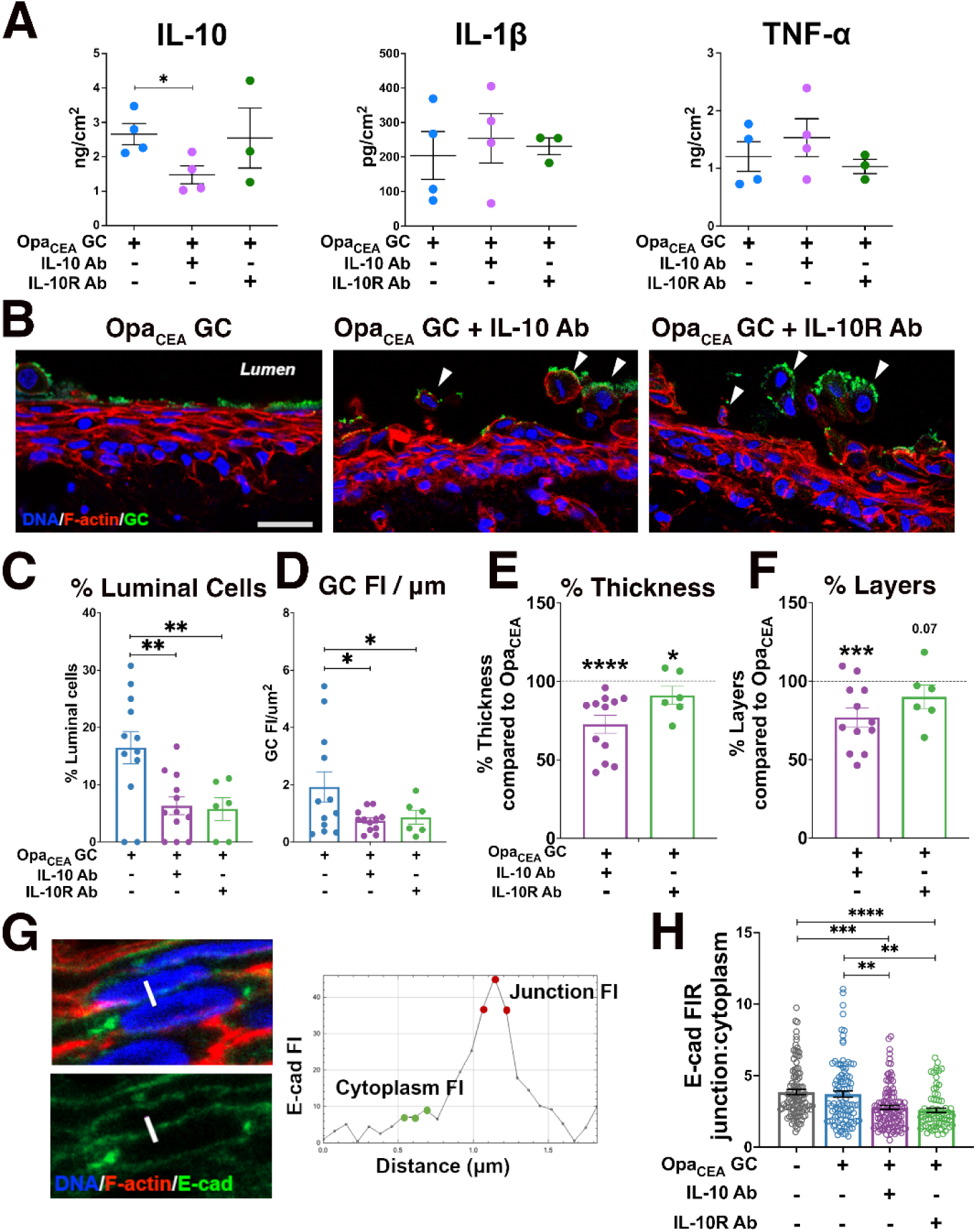
IL-10 neutralization and IL-10 receptor-blocking antibodies reduce GC colonization by increasing ectocervical epithelial cell shedding. Human cervical tissue explants were incubated without and with MS11Opa_CEA_ in the absence and presence of anti-IL-10 (10 µg/ml) or IL-10Rα antibody (5 µg/ml) for 24 h (MOI~10). The culture supernatants were collected, and tissues were cryopreserved. (**A**) The concentrations of IL-10, IL-1β, and TNF-α in the supernatants were measured by Luminex Magpix (IL-10, IL-1β, and TNF-α). Data points represent individual cervixes. n=3-4. (**B)** Representative images of ectocervical tissue sections stained for GC, DNA, and F-actin. Arrowhead, shed epithelial cells. Scale bar, 10 μm. (**C** and **D**) Quantification of GC colonization by the percentage (±SEM) of luminal epithelial cells associated with GC (**C**) and by FI (±SEM) of GC staining per μm^2^ of the luminal surface (**D**). Shown are the average values from 2-4 cervixes, 3 independent analyses per cervixes, and 4-8 randomly acquired images per analysis. Data points represent independent analysis. (**E** and **F**) Quantification of epithelium shedding by the percentages (±SEM) of remaining epithelium thickness and cell layers in tissue explants inoculated with MS11Opa_CEA_ in the presence of IL-10 or IL-10Rα antibody, compared to tissue explants inoculated with MS11Opa_CEA_ without IL-10 or IL-10Rα antibody. Shown are the average values from 2-4 cervixes and 3 independent analyses per cervix. Data points represent independent analysis. (**G** and **H**) Disruption of epithelial cell-cell junctions was determined by the FI ratio (FIR) of E-cad at the cell-cell border (red dots in **G**) relative to the cytoplasm (green dots in **G**) using FI line profiles (**G**) crossing cell-cell junction (**H).** Shown are the average FIR (±SEM) from 2-4 cervixes with 3 independent analyses per cervix, 4-10 randomly acquired images per analysis, and 3 line profiles per image. Data points represent individual line profiles. \*\**p*<0.01, \*\*\**p*<0.001, \*\*\*\**p*<0.001, by unpaired student’s *t*-test.

### GC induce IL-10 production by engaging CEACAM1 and activating SHP1/2

Our previous studies have shown that GC interaction with CEACAM1 via Opa_CEA_ increases GC colonization at the ectocervix by inhibiting epithelial cell shedding. This increase depends on the ITIM motif within the cytoplasmic domain of CEACAM1 and subsequent activation of SHP1/2 (61, 62). Since MS11Opa_CEA_ but not MS11ΔOpa increase cervical secretion of IL-10, we questioned whether GC induce IL-10 production through CEACAM1. We treated ectocervical tissue explants with NSC-87877, a membrane-permeable small molecule inhibitor of SHP1/2 (63) that inhibits the effects of Opa_CEA_ on GC colonization and cervical epithelial shedding (18). We collected the supernatants from explants inoculated without or with MS11Opa_CEA_ or MS11ΔOpa in the absence or presence of NSC-87877 (20 µM) and determined the concentration of IL-10, IL-1β, TNF-α, and TGF-β using Luminex Magpix or ELISA. Treatment with the SHP1/2 inhibitor restored the secreted IL-10 level by MS11Opa_CEA_-inoculated tissue explants to uninoculated and MS11ΔOpa-inoculated tissue explants (Figure 5A). However, treatment of the SHP1/2 inhibitor did not significantly affect the secretion levels of the pro-inflammatory cytokines IL-1β (Figure 5B) and TNF-α (Figure 5C) and the pleiotropic cytokine TGF-β (Figure 5D). This result suggests that GC induce IL-10 production but not IL-1β and TNF-α through CEACAM1 and the downstream SHP1/2, and GC-induced IL-10 contributed to Opa_CEA_-mediated colonization enhancement.

**Figure 5.**
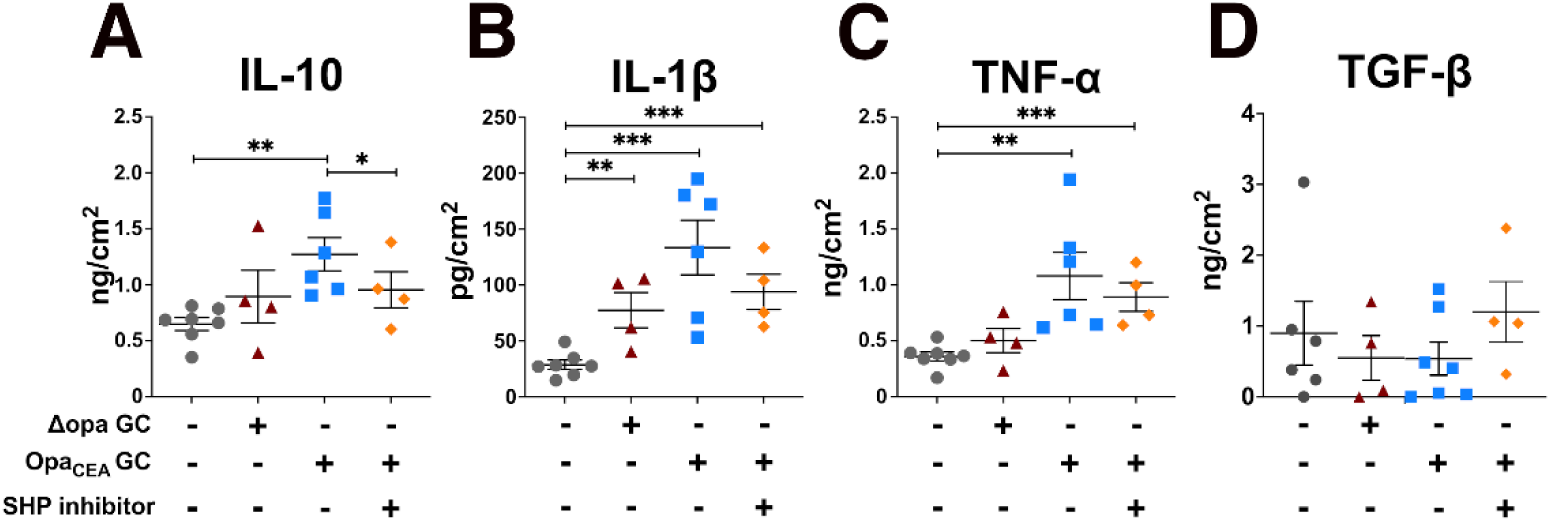
GC-induced IL-10 but not IL-1β and TNF-α secretion by human cervical tissue explants requires CEACAM1 signaling. Supernatants of cervical tissue explants inoculated without or with MS11Opa_CEA_ or ΔOpa in the absence or presence of the SHP1/2 inhibitor NSC-87877 (20 μM) for 24 h were collected. The cytokine concentrations were measured by Luminex Magpix or ELISA (TGF-β) and normalized to the luminal surface area and the supernatant volume of each explant. Shown are the average concentration (±SEM) from 4-7 cervixes. Data points represent individual cervixes. * *p*<0.05, ** *p*<0.01, *** *p*<0.001 by unpaired student’s *t*-test.

## Discussion

While the mechanisms by which GC infect the human cervical epithelium have been studied previously (11, 18, 64–66), the initial local cytokine response and its impact on GC infection remains unknown. Using human cervical tissue explants that were disconnected from lymphatic and blood circulation, we show that GC inoculation increases the secretion levels of both the pro-inflammatory cytokines IL-1β and TNF-α and the anti-inflammatory cytokines IL-10 but not TGF-β during the first 24 h. The locally secreted anti-inflammatory cytokine IL-10, but not inflammatory cytokines IL-1β and TNF-α, modulates GC infection at the human cervix. The expression of an Opa protein that binds CEACAMs is required for GC to increase the local cytokine production. GC increase the secretion of IL-1β and TNF-α, but not IL-10, by activating NF-κB in both cervical epithelial and subepithelial cells. GC elevate the secretion of IL-10 but not IL-1β and TNF-α by binding to CEACAMs and activating their downstream SHP1/2. GC-induced IL-10 enhances GC colonization by inhibiting epithelial cell-cell junction disassembly and GC-associated cervical epithelial cell shedding. Reductions of IL-1β and TNF-α secretion by inhibiting NF-κB did not affect GC infection.

A significant finding of this study is that during the first 24 h, GC colonization elevates the local secretion of the anti-inflammatory cytokine IL-10 at the cervix. While IL-10 was detected in women’s cervical mucus, its level was reported to be increased, decreased, or unchanged by GC infection (30–33). However, in these studies, how long women had been infected before sample collection and whether infected women were symptomatic or asymptomatic were unknown. GC has been shown to elevate IL-10 levels of mouse genital track explant cultures (39), even though mice do not express GC host receptors, such as human CEACAMs, and GC-targeted human immune factors, such as complement factor H (67, 68). Culturing GC with monocyte-derived dendritic cells (34) and macrophages (36), CD4+ T cells (37), and mononuclear cells (38) from human peripheral blood all increases IL-10 production. However, during the initial stages of natural infection (less than 24 h after exposure), GC primarily colonize the ectocervical epithelium and have little chance of encountering subepithelial immune cells. Since the human cervical tissue explants are surgically disconnected from the blood and lymphatic circulation, our data provide the first evidence for GC-elevated mucosal secretion of IL-10 by human cervical cells. These data together demonstrate the robust ability of GC to induce the production of IL-10, a potent anti-inflammatory cytokine, by the local mucosal and central immunity despite the types of cells they encounter. The local elevation of IL-10 at the cervix during the first 24 h of infection can potentially prevent inflammation, thus enabling GC colonization without symptoms.

This study found that cervical tissue explants constitutively secreted IL-10 at levels relatively higher than the basal level of the inflammatory cytokines IL-1β and TNF-α. While tissue wounding caused by surgery, tissue handling, and initial antibiotic treatment could modulate cytokine production, the relatively high basal level of IL-10 suggests an anti-inflammatory cytokine environment of the human cervix. The anti-inflammatory environment is consistent with the physiological functions of the cervix, a gate between the vagina and the uterus, which requires the cervix to tolerate semen and microbiota. The FRT is known to upregulate IL-10 production during pregnancy, which is critical for the immune tolerance of fetuses (69, 70). Our finding that inhibiting NF-κB, leading to a reduction in inflammatory cytokines, did not impact GC colonization suggests a prevailing anti-inflammatory cytokine-dominant environment in the human cervix. GC-induced IL-10 secretion further enhances this anti-inflammatory cytokine environment. In addition to IL-10, other locally secreted pleiotropic cytokines, such as leukemia inhibitory factor (LIF) and TGF-β, potentially contribute to the immunosuppressive environment of the cervix, and their role in asymptomatic GC infection remains to be explored.

Our results suggest that locally secreted IL-10 can directly act on cervical epithelial cells to promote GC colonization by inhibiting the disassembly of epithelial cell-cell junctions and GC-associated epithelial cell shedding. There are several potential mechanisms by which IL-10 can promote GC colonization at the cervix: activating epithelial cell proliferation, differentiation programs, and wound repair pathways by directly engaging IL-10R on epithelial cells (40, 42), inhibiting pro-inflammatory cytokine production and interfering with inflammatory cytokine-induced signaling pathways (40, 71). Inflammatory cytokines, i.e. IL-1β and TNF-α, can disrupt epithelial cell-cell junction by activating myosin and inhibiting junction protein expression by inducing cellular signaling (44, 48, 72), which leads to epithelial shedding. Here, we show that IL-10 neutralization or IL-10Rα blocking by antibodies weakens the epithelial cell-cell junction but does not increase the secretion of IL-1β and TNF-α by the cervix, suggesting that the effect of IL-10 we observed is not mediated through downregulating inflammatory cell junction-disrupting cytokines. However, locally secreted IL-10 may indeed inhibit inflammatory cytokine production *in vivo*. The unchanged levels of inflammatory cytokines we observed were likely due to the failure of IL-10 and IL-10Rα antibodies to reach inflammatory cytokine-producing cells in the subepithelium of the cervix. The primary location of IL-10Rα blocking antibody at the luminal layer of the ectocervical epithelium suggests that IL-10Rα antibody only blocked IL-10 from binding IL-10R at the luminal surface of cervical epithelial cells. IL-10-mediated inhibition of shedding GC-associated cervical epithelial cells in the presence of the potent inflammatory cytokines IL-1β and TNF-α underscores the importance of the direct interaction of IL-10 with IL-10R on epithelial cells for GC-establishing infection at the human cervix. Using the mouse vaginal infection model, Liu et al. showed that GC-induced IL-10 promotes repeat GC infection by suppressing the activation of adaptive immunity, both cellular and humoral immunity (39). If GC cervical infection could induce IL-10 production by draining lymph nodes and blood circulation, we can expect a similar IL-10-mediated suppression of adaptive immunity in humans.

The elevated levels of inflammatory cytokines and nuclear NF-κB, a crucial transcriptional factor that controls the production of inflammatory cytokines (56, 57), by GC inoculation suggest that GC cannot escape immune detection. Instead, GC increase the secretion of anti-inflammatory cytokines to counteract inflammation activation and maintain or even enhance immune tolerance in the cervix, depending on the relative levels of pro- and anti-inflammatory cytokines. Cervical cells can detect GC through Toll-like receptors (TLRs) and inflammasomes (23, 73). Interestingly, MS11Opa_CEA_ that primarily colonize the luminal surface of the cervical epithelial cells activate NF-κB in both epithelial and subepithelial cells, suggesting the activation of NF-κB in subepithelial cells may be indirect, probably through inflammatory cytokines secreted by cervical epithelial cells. Surprisingly, MS11ΔOpa, which can penetrate into the subepithelium of the transformation zone and the endocervix and have the possibility of directly interacting with subepithelial immune cells, induce a lower level of an increase in IL-1β than MS11Opa_CEA_ and no elevation of TNF-α and IL-10. The weak local cytokine response induced MS11ΔOpa is associated with a lower level of increases in NF-κB in the nuclei of epithelial cells and even reductions in NF-κB in the nuclei of subepithelial cells. As MS11Opa_CEA_ was generated by introducing an Opa_CEA_ to MS11ΔOpa (74), our data suggest a requirement of the Opa-CEACAM interaction to induce IL-1β and TNF-α production. However, inhibition of CEACAM1 downstream SHP1/2 slightly but not significantly reduced MS11Opa_CEA_-induced IL-1β and TNF-α, which argues against this notion. Besides activating SHP1/2, Opa-CEACAM interactions enhance GC colonization efficiency at the cervix, increasing the number of GC that interact with epithelial cells directly, which potentially enables higher levels of TLR and inflammasome activation, leading to their downstream NF-κB activation and pro-inflammatory cytokine secretion. This multifaceted hypothesis remains to be tested. Since MS11ΔOpa-induced NF-κB reduction in the nuclei of the subepithelial cells, it suggests an Opa-independent mechanism for GC suppression of inflammatory responses.

This study suggests that GC elevate the IL-10 secreting levels in the human cervix by engaging CEACAM1, which contains an ITIM motif in the cytoplasmic tail and can activate SHP1/2. Our supporting data include the induction of IL-10 by MS11Opa_CEA_ but not MS11ΔOpa and reductions of both IL-10 secretion and MS11Opa_CEA_ cervical colonization (18) by the SHP1/2 inhibitor. Which types of cervical cells respond to CEACAM1 binding to secret IL-10 remains unknown. While immune cells are the primary sources of IL-10, non-immune cells, such as epithelial cells, can secrete IL-10 as well (42, 75). Besides cervical epithelial cells (18), many types of immune cells express CEACAM1 (61). CEACAM1 functions as a negative regulator in T cells by cis-homo interactions on the same cells and trans-homo interactions with other cells (76–78). CEACAM1 knockout in mice leads to the hyper-expansion of conventional T cells but the reduction of regulatory T cells, IL-10 producers, in the liver (79). *In vitro* experiments have shown that GC inhibits T cell receptor-mediated T cell activation by engaging CEACAM1 on the T cell surface by Opa and activating CEACAM1-downstream SHP1/2, downregulating T cell receptor signaling (80, 81). GC has also been shown *in vitro* to promote macrophage polarization towards an alternatively activated phenotype characterized by heightened IL-10 production (57). Even though GC Opa proteins do not bind mouse CEACAMs (82), GC-induced IL-10 production in mice lymphocytes partially depends on Opa expression (39). Its underlying mechanisms remain to be determined. Accumulating evidence supports Opa-CEACAM1 interactions as one of the mechanisms for GC to induce IL-10 production.

Our study reveals a new mechanism employed by GC to evade mucosal immunity, enhancing local secretion of the anti-inflammatory cytokine IL-10 at the mucosal surface of the cervix by engaging CEACAM1. IL-10 then directly acts on cervical epithelial cells, preventing GC-colonizing epithelial cells from shedding off, in addition to protecting GC from immune detection. This mechanism potentially enables GC to establish asymptomatic infection in women and explains why CEACAM-binding Opa proteins have been evolutionally favored (most of the 11 Opa isoforms bind CEACAMs) (21, 22) and why GC isolated from vaginally infected mice (83) and experimentally inoculated men (84) all express Opa proteins.

## Methods

### Ethics statement

Human cervical tissues were obtained through the National Disease Research Interchange (NDRI, Philadelphia, PA). Human cervical tissues used were anonymized. The use of human tissues has been approved by the Institution Review Board of the University of Maryland.

### Neisseria strains

*N. gonorrhoeae* strain MS11 with all 11 *opa* gene deleted (ΔOpa) and MS11 ΔOpa expressing non-phase variable CEACAM-binding Opa_52_ (Opa_CEA_) (74) were used. Piliated bacteria were identified based on colony morphology using a dissecting light microscope. GC were grown on plates with GC media and 1% Kellogg’s supplement (GCK) for 16–18 h before inoculation. The concentration of bacteria in suspension was determined by using a spectrophotometer.

### Human cervical tissue explants

Healthy cervical tissues were obtained from patients (28–42 years old) undergoing hysterectomies for medical reasons unrelated to the cervix and received within 24 h post-surgery through National Disease Research Interchange (NDRI). Tissue explants were generated and processed using a previously published protocol (85). Briefly, Muscle parts of the tissue were removed using a carbon steel surgical blade. Cervical tissues from one human subject were cut into three or four equal pieces with the dimension of ~2.5 cm (L) X 0.6 cm (W) X 0.3 cm (H) for comparing different inoculation conditions. For ectocervix tissue explants, ectocervical tissues were separated from the transformational zone and endocervix by visual inspection. Tissue explants were incubated in the CMRL-1066 (11530037, Gibco), containing 5% heat-inactivated fetal bovine serum (A5256701, Gibco), L-glutamine (2 mM, 25030081, Gibco), bovine insulin (1 μg/ml, 16634, Sigma-Aldrich), and penicillin/streptomycin for 24 h, followed by antibiotic-free media for another 24 h. GC were inoculated at MOI ~10 (10 bacteria to 1 luminal epithelial cell). The number of epithelial cells at the luminal surface was determined by the luminal surface area of individual explants divided by the average luminal area of individual cervical epithelial cells (25 μm^2^). The cervical tissue explants were incubated with GC at 37°C with 5% CO_2_ with gentle shaking for 24 h and washed with antibiotic-free cervical tissue culture medium at 6 and 12 h to remove non-adhered bacteria.

### Inhibitor and antibody treatment

Cervical tissue explants were incubated with bacteria in the presence or absence of NF-κB inhibitor BAY 11-7082 (3 μM, HY-13453, MedChemExpress) (58), SHP inhibitor NSC-87877 (86) (20 μM, 565851-50MG, EMD Millipore), anti-human IL-10 antibody (10 μg/ml, 10100-01, SouthernBiotech), or anti-human IL-10Rα antibody (5 μg/ml, 308802, BioLegend) for 24 h. The inhibitors and the antibodies were replenished after the 6 and 12 h washing.

### Quantification of Cytokine secretion

*S*upernatants were collected from cervical tissue explant cultures at 6, 12, and 24 h post-inoculation, pooled together in an 1:1:2 volume ratio, and a protease inhibitor cocktail (PIC0002, Sigma-Aldrich) was added to prevent protein degradation. Cytokines in the supernatants were quantified using a Luminex Magpix system (Bio-Plex® MAGPIX™ Multiplex Reader, Bio-Rad Laboratories) and the Human XL Cytokine Luminex Performance Assay 46-Plex Fixed Panel (Bio-Techne), according to the manufacturer’s protocol. The secretion levels of IL-1β and TGF-β were quantified using enzyme-linked immunosorbent assay (ELISA) by human IL-1β ELISA Max Deluxe kit (437004, BioLegend) and human TGF-β1 duoset ELISA kit (Dy204-05, R&D System). Each sample was run in duplicate. The cytokine concentrations of each sample by either Luminex Magpix or ELISA were normalized based on the supernatant volume and the tissue luminal surface area.

### Immunofluorescence analysis of human cervical tissue explants

The tissue explants were fixed in 4% paraformaldehyde 24 h post-inoculation, embedded in 20% gelatin, cryopreserved, sectioned crossing the luminal, basal surfaces of the epithelium and subepithelial tissues, stained for F-actin (100 nM, phalloidin, PHDH1, Cytoskeleton), E-cadherin (5 µg/ml, 610182, BD Bioscience), NF-κB p65 (5 µg/ml, 8242, Cell Signaling Technology), and GC by specific antibodies, and nuclei by Hoechst (20 µg/ml, H3570, Life Technologies), and imaged using confocal fluorescence microscope (CFM, Zeiss LSM 980, Carl Zeiss Microscopy LLC). Images were acquired using Zeiss Zen software.

The levels of NF-κB activation were quantified by the mean fluorescence intensity (MFI) of p65 staining in individual nuclei. Individual nucleus was identified by Hoechst staining using NIH ImageJ. The data were generated using 3-4 human cervixes, 2 independent analyses per cervix, 5-11 randomly acquired images per ectocervical region, 3-8 randomly acquired images per TZ region, 3-10 randomly acquired images per endocervical region per analysis, including 200 nuclei per ectocervical epithelial region, 70 nuclei per ectocervical subepithelial region, 35 nuclei per transformation zone epithelial region, 45 nuclei per transformation zone subepithelial region, 35 nuclei per endocervical epithelial region, and 170 nuclei per endocervical subepithelial region per analysis.

The levels of GC colonization at the ectocervix were quantified by two methods using CFM images (18): (1) the percentage of luminal epithelial cells with GC attached at the luminal surface by manually accounting and (2) the fluorescence intensity (FI) of GC staining per μm^2^ of the luminal surface using the NIH ImageJ software. The data were generated from 2-4 human cervixes, 3 independent analyses per cervix, and 3-10 randomly acquired images per analysis.

The levels of epithelial shedding in the ectocervix were determined by two methods (18): (1) the percentage of the remaining thickness (μm) of the epithelium and (2) the percentage of remaining epithelial cell layers (based on F-actin staining) compared to uninfected ectocervical tissue explants, using the NIH ImageJ software. The data were generated from 2-4 human cervixes, 3 independent analyses per cervix, and 3-10 randomly acquired images per analysis.

The redistribution of E-cadherin from the cell-cell junction to the cytoplasm was evaluated by the fluorescence intensity ratios (FIR) of E-cadherin staining at the cell-cell junction relative to the cytoplasm in individual epithelial cells using CFM images by the NIH ImageJ software. The data were generated from 2-4 human cervixes, 3 independent analyses per cervix, and 4-10 randomly acquired images per analysis.

### Gene expression analysis by NanoString

At 24 h post-inoculation, the tissue explants were embedded in Tissue-Tek® O.C.T. Compound (4583, Sakura Finetek) and snap froze in liquid nitrogen. Thirty 10-μm thickness tissue sections were collected from each tissue explant for RNA isolation using RNeasy Mini Kit (74104, Qiagen). The multiplexed NanoString nCounter™ Immunology Panel (PSTD Hs Immunology V2-12, NanoString Technologies) with 594 genes was used, according to the manufacturer’s protocol. The data QC, background threshold, normalization, and differential gene expression analysis were conducted using nSolver™ Analysis software. The quality of the run for each sample was confirmed by the quality control, including the 6 spiked-in RNA positive controls and the 8 negative controls present in the panel, the FOV (fields of view per sample) counted, and the binding density. Gene expression data were normalized by using the 15 housekeeping genes present in the panels.

Background level was determined by mean counts of 8 negative control probes plus two standard deviations. When a gene’s raw counts are below the background of >50% of all samples, this gene is excluded from differential expression analysis. Differential gene expression analysis was performed by comparing the normalized count of each gene in infected samples and uninfected samples, and genes with *p* < 0.05 were identified as differential expression genes.

### Statistical analysis

Statistical significance was assessed using the Student’s *t*-test by Prism software (GraphPad). P values were determined by unpaired t-test with Welch’s correction.

## Acknowledgments

We thank the UMD CBMG Imaging Core for all microscopy experiments and the Maryland Genomics Core at the University of Maryland School of Medicine for the NanoString transcriptomic analysis. This work was supported by a grant from the National Institutes of Health to DCS and WS, AI123340.

