## Supplementary Figures for "Neisseria gonorrhoeae induces local secretion of IL-10 at the human cervix to promote asymptomatic colonization"

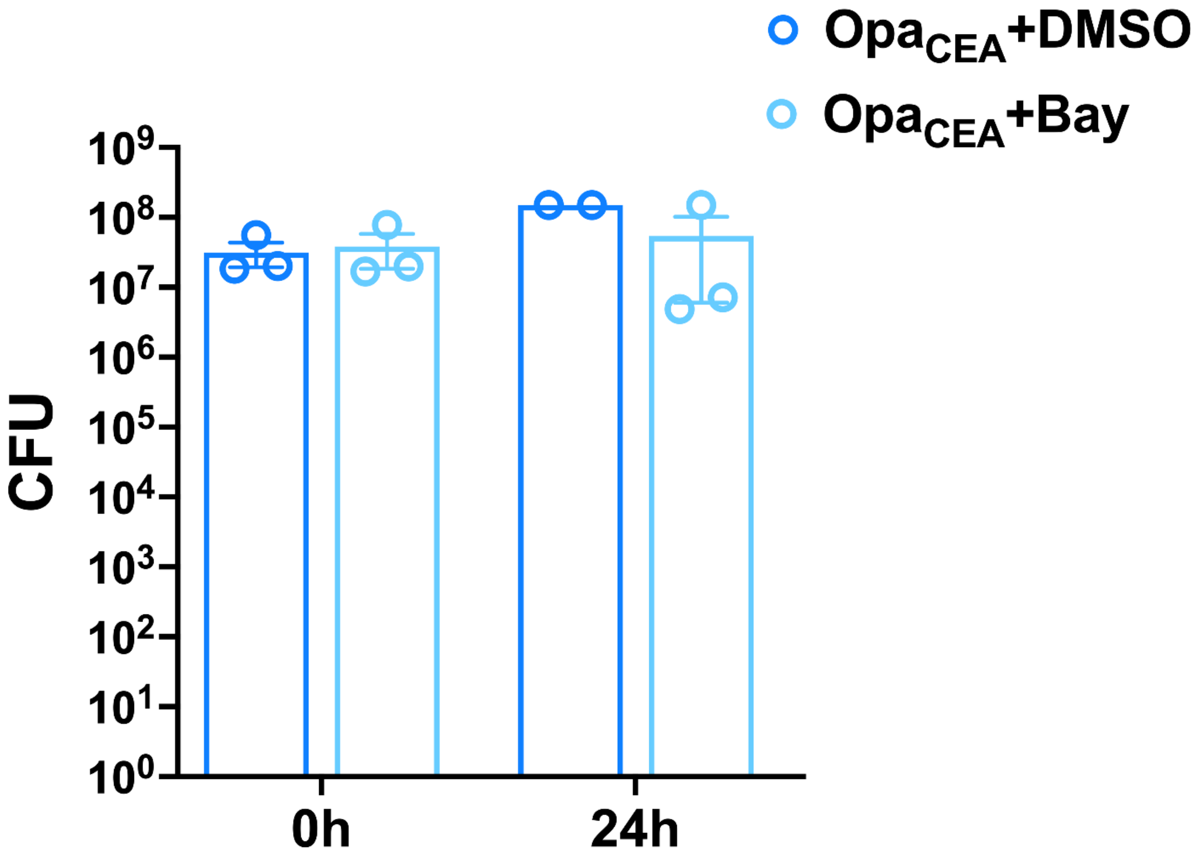


**Supplementary Figure 1. Treatment of the NF-κB inhibitor Bay 11-7082 does not affect GC growth.** Human ectocervical tissue explants were inoculated with MS11OpaCEA (MOI~10) in the absence and presence of the NF-κB inhibitor Bay11-7082 (3 μM) for 24 h. Bacteria were enumerated by counting CFU after serially diluting tissue culture media and plating on GCK plates. Shown are the averages of 2-3 independent experiments.

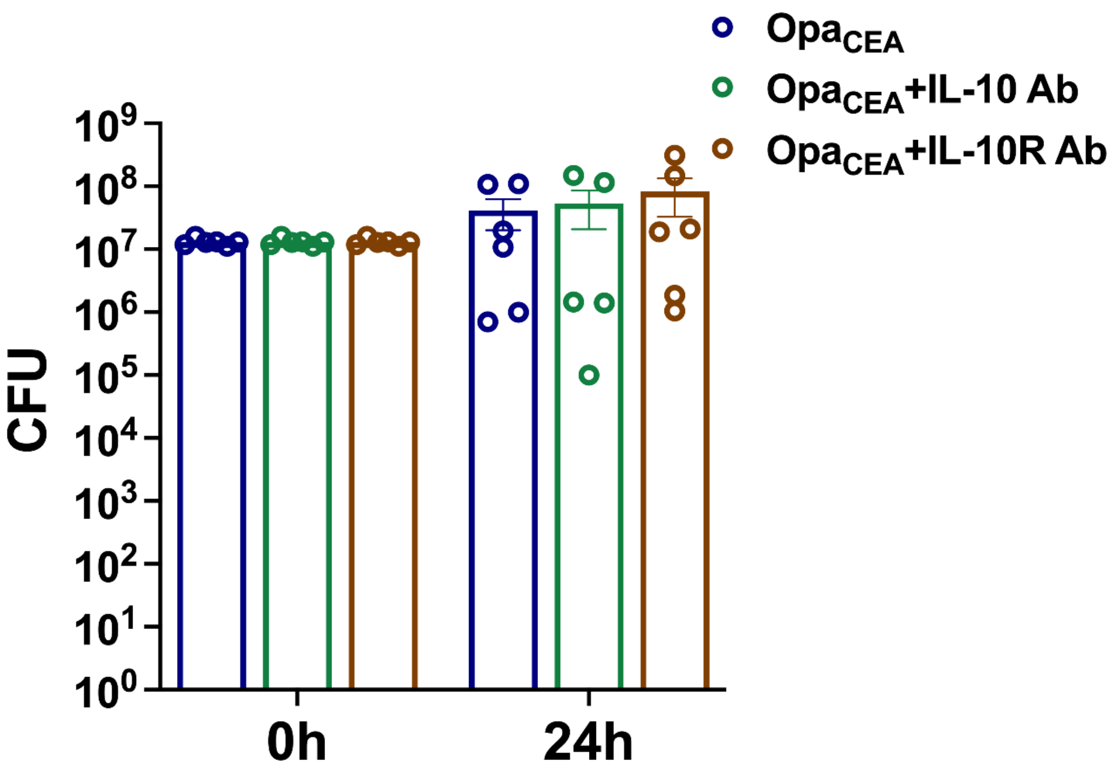


**Supplementary Figure 2. IL-10 neutralization and IL-10 receptor α-blocking antibodies do not affect GC growth.** MS11Opa$_{CEA}$ were cultured in CMRL-1066 media containing 5% FBS for 24 h in the absence or presence of the IL-10 neutralization (10 μg/ml) and IL-10 receptor α-blocking antibodies (5 μg/ml). Bacteria were enumerated by counting CFU after serially diluting tissue culture media and plating on GCK plates. Data points represent individual wells. Shown are the averages of 3 independent experiments with two wells per experiment.

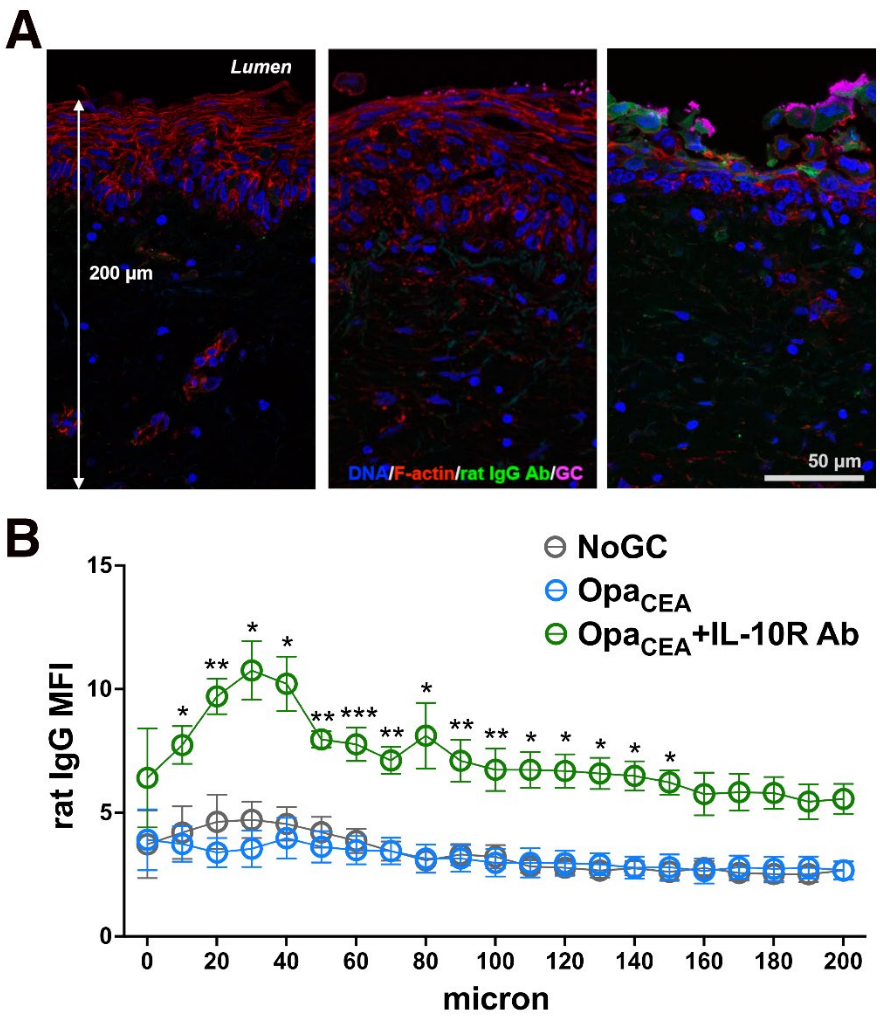


**Supplementary Figure 3. IL-10Rα-blocking antibody primarily targets cervical epithelial cells.** Human ectocervical tissue explants were incubated without and with MS11Opa$_{CEA}$ (MOI~10) in the absence and presence of rat IgG anti-human IL-10Rα antibody (5 µg/ml) for 24 h and cryopreserved. Tissue sections were stained with anti-rat-IgG, GC-specific antibody, phalloidin, and Hoechst and imaged by CFM. (**A**) Representative images of the ectocervical tissues. Scale bar, 50 µm. (**B**) Quantification of rat IgG MFI from the luminal surface of the ectocervical epithelium to 200 µm depth into the subepithelium. Shown are the average values (±SEM) generated from 3-4 ectocervixes and 5-8 randomly taken images per cervix.
